# ECD of PepT1 interacts with TM1 to facilitate substrate transport

**DOI:** 10.1101/2022.03.01.482558

**Authors:** Jiemin Shen, Miaohui Hu, Zhenning Ren, Xiao Fan, Corinne Portioli, Xiuwen Yan, Mingqiang Rong, Ming Zhou

## Abstract

Mammalian peptide transporters, PepT1 and PepT2, mediate uptake of a wide variety of di- and tri-peptides and are essential for the absorption of dietary peptides in the digestive tract and the recovery of peptides in renal filtrate. PepT also mediates absorption of many drugs and prodrugs to enhance their bioavailability. PepT has 12 transmembrane (TM) helices that fold into two domains, the N-terminal domain (NTD, TM1-6) and C-terminal domain (CTD, TM7-12), and a large extracellular domain (ECD) of ∼200 amino acids between TM9 and TM10. It is known that peptide transport involves large motions of the N- and C-domains, but the role of ECD remains unclear. Here we report the structure of PepT1 from *Equus caballus* (horse) determined by cryo-electron microscopy. The structure shows that ECD interacts with TM1 and bridges the N- and C-domains. Deletion of the ECD or mutations to the TM1-ECD interface both impair the transport activity. These results demonstrate a role of ECD in structure and function of PepT1 and enhance our understanding of the mechanism of transport in PepT1.

## Introduction

Mammalian peptide transporters, PepT1 and PepT2, are members of the solute carrier (SLC) transporter family 15^1^. PepT1 is mainly found on intestinal brush border membranes and mediates uptake of small peptides^2,3^. PepT2 is found in epithelial cells in kidney and retrieves peptides from the glomerulus filtrate^1,4,5^. Due to their broad substrate spectrum, PepT1 and PepT2 also transport a large variety of small-molecule drugs such as antibiotics and antiviral drugs, and prodrugs with amino acids as part of the scaffold^6^. The concentrative uptake of peptides or small-molecule drugs by PepT is achieved by co-transport of protons and is thermodynamically driven by both the pH gradient and the membrane potentials^1^.

Structures of human PepT1 and PepT2^7^, rat PepT2^8^, and several bacterial homologs^9-20^ have been reported previously. The structures show that PepTs have 12 transmembrane helices (TM1-12) which fold into two well-defined domains. TM1-6 form the N-terminal domain (NTD), and TM6-12 the C-terminal domain (CTD). A peptide binding site is located approximately in the middle of membrane and lined by residues from TM1, 2, 4, and 5 of the NTD and TM7, 8, and 10 of the CTD. PepT structures have also been captured in conformations with the peptide binding site being solvent accessible to either the extracellular side, the outward-facing conformation, or the intracellular side, the inward-facing conformation, and a few conformations in between^7,8,21^. The structures show that the transition between the inward- and outward-facing conformations comes from rigid-body motions of the NTD and CTD. This is often referred to as the rock-switch model of alternating access, and is shared by other transporters of the major facilitator superfamily (MFS) fold^22^.

Different from bacterial homologs, mammalian PepT1 and PepT2 have an extracellular domain (ECD) composed of ∼200 residues located between TM9 and TM10. The molecular weight of ECD is approaching to that the NTD or CTD. The ECD has a well-defined structure composed mainly of β-strands folded into two immunoglobin-like subdomains^7,8,23^. The structure of human PepT2 captured in the inward-facing conformation (PDB ID: 7PMY) shows that the ECD is in close proximity to the NTD, but no specific interactions were identified^7^. A previous mutational study showed that the PepT2 function is not perturbed by deletion or swapping of its ECD with that of PepT1^23^, however, the role of ECD in PepT1 was not assessed because PepT1 with its ECD deleted or swapped with that of PepT2 do not have sufficient level of expression in the *Xenopus* oocytes system^23^.

In this study, we determined the structure of PepT1 from horse (*Equus caballus*) in the inward-facing conformation. The structure shows that its ECD interacts with the NTD, which prompted us to examine the function of ECD in PepT1. We measured the transport activities of PepT1 in human embryonic kidney (HEK) cells, and found that deletion of the ECD or mutations to the interface of ECD and NTD significantly reduce the rate of substrate transport. These results led us to conclude that interactions between ECD and NTD are functionally relevant and may facilitate transition of PepT1 from the outward-facing to the inward-facing conformation.

## Results

### Proton-coupled peptide transport by horse PepT1

To examine transport activity of horse PepT1, we expressed it in HEK293 cells and measured uptake of peptide and proton. As shown in **Fig. 1a**, cells expressing horse PepT1 accumulate a labeled dipeptide (^3^H-Ala-Ala) while cells transfected with a control vector do not accumulate significant amount of the peptide (**Fig. 1a**). We also measured co-transport of protons with peptide substrates by first loading the cells with a pH sensitive dye and monitoring intracellular pH changes. Proton uptake occurs in the presence of the dipeptide substrate (Ala-Ala, 1mM), and lower the external pH, higher the amount of proton uptake (**Fig. 1b** & **1c**, and **Extended Data Fig. 1**). As a further test that the observed proton uptake is induced by peptide transport, we measured pH changes in the presence of two additional di-peptides, Glu-Glu, and Gly-Sar, and found that although all three dipetides are transported, Ala-Ala inducing the largest fluorescence increase (**Extended Data Fig. 1d**). The expression level of PepT1 between different batches of transfection was estimated by Western blot (Methods and **Extended Data Fig. 2**) and the amount dipeptide or proton uptake was normalized accordingly. These results indicate that horse PepT1 is a proton coupled peptide symporter similar to human PepT1^24^.

**Fig. 1:**
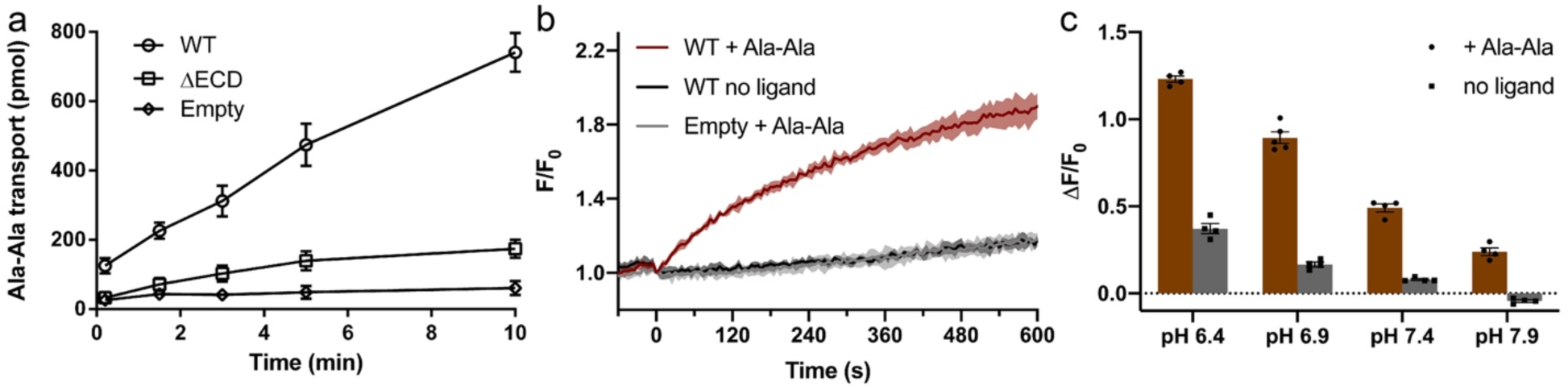
Functional characterization of horse PepT1 with cell-based assays. **a**, Peptide uptake activity of horse PepT1 WT and ΔECD in HEK293 cells. Error bars represent standard error of the mean (SEM) from three repeats. **b**, pH changes induced by proton-coupled peptide uptake. Ligand Ala-Ala (red) or blank buffer (dark gray) was added to HEK293 cells expressing horse PepT WT. Cells transfected with empty vector were added with the same ligand (light gray). The fluorescence changes (F/F_0_) of a pH sensitive dye were used to indicate cytosolic pH changes. The time courses are shown as solid line (mean) with shaded region (standard deviation, SD) from four repeats. **c**, Fluorescence changes after 10 min of transport in the outside buffer of different pHs. Two-way ANOVA: among different pHs, p < 0.0001; with or without ligand, p < 0.0001; interaction, p < 0.0001. For all bar graphs in this paper, a scatter plot is overlaid on each bar and the height is the mean of at least three repeats with the error bar representing the SEM.

### Cryo-EM structure of horse PepT1 in nanodisc

We expressed and purified the horse PepT1 from Sf9 (*Spodoptera frugiperda*) cells and reconstituted the protein into lipid nanodiscs for structure determination by cryo-electron microscopy (**Fig. 2 and Methods**). The elution volume of horse PepT1 from a size-exclusion chromatography column is consistent with it being a monomer (**Fig. 2a**). PepTs of human and rat origins were also shown to exist as a monomer^7,8^. The amino acid sequence of horse PepT1 is 84% identical and 96% similar to that of human PepT1. The soluble ECD served as a fiducial marker for particle alignment during 3D reconstruction, and we were able to obtain a cryo-EM map with an overall resolution of ∼3.6 Å (**Fig 2b**, and **Extended Data Fig. 3**). The map for the TM domain, which is mainly composed of α-helices, is of sufficient quality for *de novo* model building. Although the map for the ECD, which is mainly composed of β-strands and not constrained by nanodisc, has modest resolution, the two subdomains and individual β-strands are readily recognizable. The overall structure of horse PepT1 is shown in **Fig. 2c**, and the fitting of individual TMs and ECD to the density map is shown in **Extended Data Fig. 4**.

**Fig. 2:**
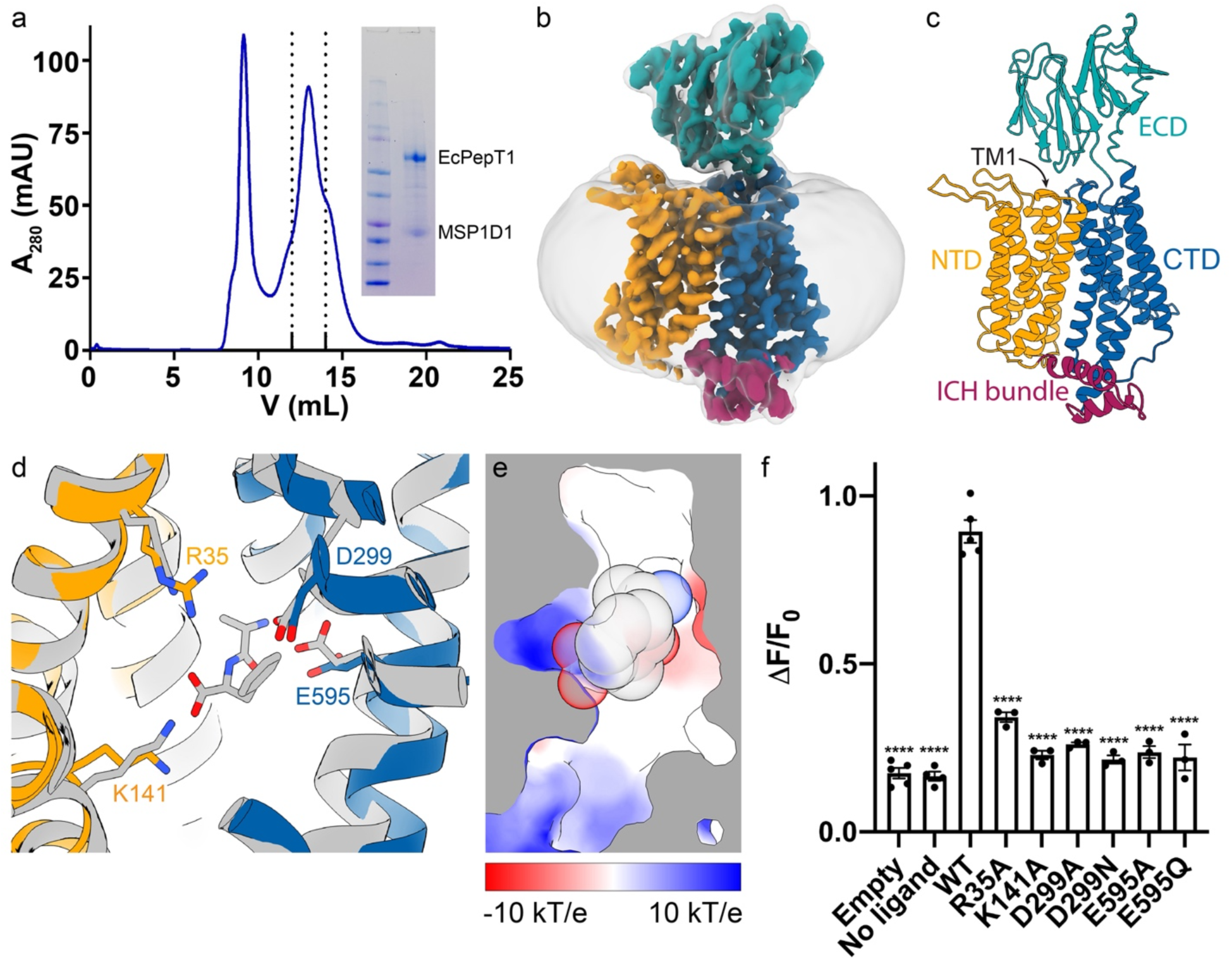
Cryo-EM structure and ligand binding site of horse PepT1. **a**, SEC profile and SDS-PAGE gel image (inset) show the reconstitution of horse PepT1 into MSP1D1 nanodisc. Fractions within the dotted lines were collected. **b**, Cryo-EM map of horse PepT1 in nanodisc colored as described in **c**. A gaussian smoothed map (translucent gray) is overlaid to display the contour of nanodisc density around the transporter. **c**, Structure of horse PepT1 in cartoon representation shows the inward-open conformation with NTD in orange, intracellular domain helix (ICH) bundle in red, and CTD in blue. **d**, Structural alignment of horse PepT1 with human PepT2 bound with an Ala-Phe ligand (PDB ID: 7PMY) colored in gray. Some conserved charged residues in the ligand binding pocket are shown as stick. **e**, Cut-through of the electrostatic surface in the binding pocket of horse PepT1 show the opposite charge on the NTD (positive) and CTD (negative). The ligand Ala-Phe shown in sphere are superimposed. **f**. Functional characterization of mutations in the ligand binding pocket. The Dunnett’s test was used as a post hoc test following one-way analysis of variance (ANOVA) with the WT group as control. Statistical significances are indicated: ****, p < 0.0001.

### Comparison of horse PepT1 to existing PepT structures

The horse PepT1 is captured in the inward-facing conformation, with the extracellular sides of its NTD and CTD making contacts and leaving the substrate binding site solvent accessible from the intracellular side. The overall structure of horse PepT1 aligns well with an inward-facing human PepT2 structure (PDB ID: 7PMY)^7^ with an overall root-mean-square deviation (RMSD) of 2.5 Å (**Extended Data Fig 5a**).

The NTD or CTD of horse PepT1 aligns well to the NTD or CTD of mammalian PepTs of known structures^7,8^ with RMSD of 1.0 to 1.4 Å. TM1-2 and TM5 in the NTD (**Extended Data Fig 5b**), and the TM7 and TM11 in the CTD (**Extended Data Fig 5c**) show larger deviations in the alignments, suggesting that these structural elements are more dynamic. The structure fold of ECD in horse PepT1 is similar that of human PepT1, PepT2, or rat PepT2 with an RMSD of 1.1 to 2.5 Å, however, the relative position of ECD to CTD varies in these structures as shown in **Extended Data Fig 5d** with the CTDs aligned. While the ECD of horse PepT1 differs from those of human PepT1 and PepT2 by less than 30° rotation, the ECD in rat PepT2 seems to be an outlier (∼79° rotation). The large change of ECD in rat PepT2 is likely caused by the presence of a nanobody used to facilitate structure determination^8^. These results suggest that the linkage between ECD and CTD is not rigid and allows flexibility for independent movement of the two domains.

The substrate binding pocket of horse PepT1 is similar to that of human PepT2^7^, and composed of conserved positive residues, Arg35 and Lys141, from the NTD, and negative residues, Asp299 and Glu595, from the CTD (**Fig. 2d**). The substrate is clamped in a cavity of opposite electric charges with its N-terminus facing negative charged surface in the CTD and C-terminus facing the positively charged surface in the NTD (**Fig. 2e**).

We mutated residues that line the substrate binding site in horse PepT1, and measured their transport activity (**Fig. 2f** and **Extended Data Fig. 2**). Significant reduction in transport activity was observed for all the mutations, and this result is consistent with similar experiments in previous reports^25,26^. These results highlight the importance of these conserved charged residues in the recognition of N- and C-terminal backbone groups on substrates. The conserved interactions with the backbone of substrates afford a conserved binding mode that is less sensitive to the side chain identity, which serves as a structural basis of the extreme substrate promiscuity of PepT1.

### Extracellular gates in PepT1

We next examined interactions between the extracellular sides of NTD and CTD, which serve as a gate to limit access of the substrate binding site to the external side. NTD and CTD make contact in three places. Asn51 in TM2 interacts with the Arg304 in TM7 (**Extended Data Fig 6a**). His58 in TM2 interacts with Asn630 in TM11, and the latter also interact with Asp299 in the substrate binding pocket (**Extended Data Fig 6b**). Arg186 at the end of TM5 interacts with Gln322 and Asp324 in the loop connecting TM7 and TM8 (**Extended Data Fig 6c**). Some of these interactions are known to be important for peptide transport, for example, His58 was implicated in mediating proton coupled transport in human PepT1^15^. Similar interactions are also observed in the structure of human PepT2 in the inward-facing conformation^7^.

The structure of horse PepT1 reveals interactions between ECD and NTD that was not observed in the structure of human PepT2 inward-facing conformation. Lys483 on the ECD interacts with the backbone carbonyl oxygen atoms of Leu45 and Phe46 at the C-terminal end of TM1 (**Fig. 3a**). The side chain of Lys483 is well-resolved in the density map. In addition, the positively charged Lys483 is further stabilized by a cation-π interaction with Phe45 on TM1 and by the helical dipole moment of TM1. Such ECD-NTD interactions do not exist in human PepT2 as the equivalent residue to the Lys in PepT1 is an Asn of no net charge on the side chain (**Extended Data Fig. 7**). Structural analysis on the extracellular side of TM1 of human PepT2^7^ shows that a histidine (His76) on the TM1-TM2 loop caps the end of TM1 (**Fig. 3b**), while equivalent residue in PepT1 is a glycine (Gly47).

**Fig. 3:**
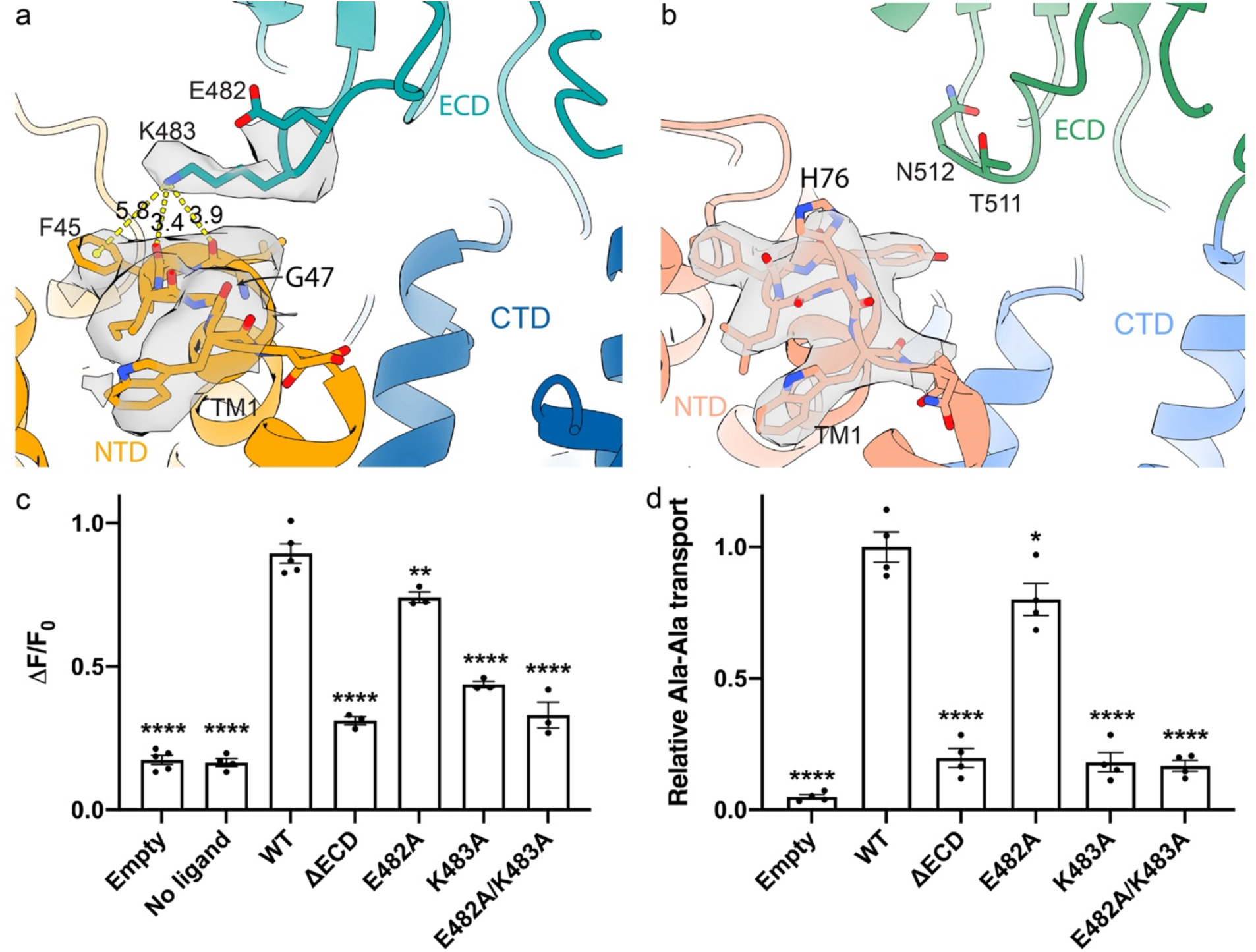
Interactions between the ECD and NTD in horse PepT1. **a**, The structure of horse PepT1 shows the ionic interactions between ECD and NTD mediated by K483. The E482, K483 and residues on TM1 and TM2 near the extracellular side are shown as stick with surrounding electron density shown as translucent gray surface. Distance measurements are indicated as yellow dash lines with labeled distances in Å. **b**, The structure of human PepT2 (PDB ID: 7PMY) shows the interactions of H76 with backbones on the C-terminus of TM1. The electron density around H76 is shown. Transport activities of horse PepT1 with mutations that affect the ECD-NTD interactions measured in the cell-based (**c**) proton transport assay and (**d**) peptide uptake assay. The Dunnett’s test was used as a post hoc test following one-way ANOVA with the WT group as control. Statistical significances are indicated: *, p < 0.05; **, p < 0.01; ****, p < 0.0001.

We then examined functional impact of the interactions between ECD and NTD. We first constructed a mutant horse PepT1 with its ECD deleted (ΔECD). ΔECD can be expressed in HEK cells and mediates proton and peptide transport, however, the rate of transport is significantly lower than these of the wild type (**Fig. 1a** and **Fig. 3c-d**). We then mutated Lys483 to Ala, and found that Lys483Ala has significantly reduced transport activity (**Fig. 3c-d**). As a control, we made an alanine mutation to Glu482 which is close to Lys483 but does not interact with TM1. Glu482Ala has a slightly reduced rate of transport compared to that of the wild type, but is substantially faster than that of Lys483Ala or the ΔECD. These results indicate that the interactions between ECD and NTD facilitate peptide transport. Further studies are required to understand how the interactions change the energy landscape of substrate transport in PepT1.

## Discussion

The horse PepT1 is the first mammalian PepT1 captured in the inward-facing conformation. The structure identified interactions between ECD and NTD, and further studies showed that the interaction has a significant functional impact. K483, which is a key residue on ECD involved in the interaction, is highly conserved in PepT1 but not in mammalian PepT2 (**Extended Data Fig. 7**). This may explain why the interaction was not observed in the structure of human PepT2 captured in the inward-facing state^7^.

The ECD-NTD interactions in PepT1 likely contribute to different transport kinetics of PepT1 and PepT2. It is well documented that the PepT1 has higher transport capacity, i.e., higher rate of transport, than PepT2^27^. The stoichiometry of the coupled protons is also different in PepT1 and PepT2^24,28^. The PepT1 was reported to require less proton to complete a transport cycle^28^. Since the substrate binding pocket of PepT1 and PepT2 are highly conserved^7^, differences in the transport kinetics may arise from the extra ECD-NTD interactions unique to PepT1. Consistent with this notion, removal of ECD in the human PepT2 resulted in no significant changes in substrate transport^23^, while our studies showed that deletion of ECD in PepT1 reduces the rate of transport. Further analysis will help pinpoint the contribution of ECD to the function of PepT1.

## Methods

### Cloning, expression, and purification of horse PepT1

The PepT1 gene from horse (*Equus caballus*, UniProt ID: <u>F6SG69</u>) was codon-optimized and cloned into a pFastBac dual vector for production of baculovirus by the Bac-to-Bac method (Invitrogen). Sf9 cells (Invitrogen/Thermo Fisher) at a density of ∼3×10^6^ cells/ml were infected with baculovirus and grown at 27 °C for 60-70 hour before harvesting. Cell pellets were frozen in liquid nitrogen and stored in −80°C fridge before purification.

Cell pellets were thawed and homogenized in lysis buffer (20 mM HEPES, pH 7.5, 150 mM NaCl, 10% glycerol) and 2mM β-mercaptoethanol, and then solubilized with 1% (w/v) Lauryl Maltose Neopentyl Glycol (LMNG, Anatrace) at 4 °C for 2 h. After centrifugation (55,000g, 45min, 4 °C), supernatants were applied to pre-equilibrated cobalt-based affinity resins (Talon, Clontech) and allowed to bind at 4 °C for 1 h under gentle rotation. The His_6_-tag was cleaved by TEV protease at room temperature for 1 hour. The flow-through containing the target protein was concentrated (Amicon 50 kDa cutoff, Millipore) and loaded onto a size-exclusion column (SRT-3C SEC-300, Sepax Technologies, Inc.) equilibrated with the FPLC buffer (20 mM HEPES, pH7.5, 150 mM NaCl) plus 1 mM (w/v) n-dodecyl-β-D-maltoside (DDM, Anatrace).

### Nanodisc reconstitution

MSP1D1 was expressed and purified following as described before^29^. A mixture of 1-palmitoyl-2-oleoyl-sn-glycero-3-phospho-(1’-rac)-choline (POPC, Avanti Polar Lipids), 1-palmitoyl-2-oleoyl-sn-glycero-3-phospho-(1’-rac)-ethanolamine (POPE, Avanti Polar Lipids) and 1-palmitoyl-2-oleoyl-sn-glycero-3-phospho-(1’-rac)-glycerol (POPG, Avanti Polar Lipids) at a molar ratio of 3:1:1, were dried and resuspended in the FPLC buffer plus 14 mM DDM^30^ by sonication. Horse PepT1, MSP1D1 and the lipid mixture were mixed at a molar ratio of 1:(2.5):(62.5) and incubated on ice for 1 hour^31^. Detergents were removed by incubation with Biobeads SM2 (Bio-Rad) overnight at 4 °C. The sample was loaded onto a size-exclusion column equilibrated with the FPLC buffer. The purified nanodisc sample was assessed by SDS-PAGE and concentrated for grid preparation.

### Cryo-EM sample preparation and data collection

Quantifoil R1.2/1.3 Cu grids were applied with 3.5 μL of purified horse PepT nanodisc at concentration of 5 or 10 mg/mL after glow-discharged with air for 15 seconds. The grids were plunged and frozen into liquid ethane cooled by liquid nitrogen using Vitrobot Mark IV (Thermo Fisher) after blotted 6 second at condition of 8°C 100% humidity. 2421 micrograph stacks in total were automatically collected with SerialEM^32^ on Titan Krios at 300 kV equipped with a K2 Summit direct electron detector (Gatan), a Quantum energy filter (Gatan) and Cs corrector (Thermo Fisher), at a nominal magnification of 105,000× with defocus values from −2.1 μm to −1.9 μm. Each stack was exposed in super-resolution mode for 5.6 s with an exposure time of 0.175 s per frame, resulting in 32 frames per stack. The total dose rate was approximately 50 e^−^/Å^2^ for each stack. The stacks were motion-corrected with MotionCor242^33^ and binned twofold, resulting in a pixel size of 1.114 Å per pixel, meanwhile dose weighting was performed^34^. The defocus values were estimated with Gctf^34^.

### Cryo-EM data processing

2,531 super-resolution movies were motion-corrected in Relion^35-37^ with 2-fold binning to generate dose-weighted micrographs (pixel size 1.114 Å). Those micrographs were then imported to cryoSPARC^38^. The blob picker was performed to 20 randomly selected micrographs and flowed by 2D classification to generate templates for auto-picking. 3,437,234 particles were picked by template picker and extracted to perform another 2D classification for particle screening. 1,504,753 particles were selected from 2D classification and used by *ab initio* reconstruction (set to 5 classes). Three 3D references (one good class and two bad classes) generated by *ab initio* reconstruction were used to perform heterogenous refinement with the original extracted 3,437,234 particles. 833,983 particles in good class were transfer to Relion^35-37^ (by csparc2star.py in pyem package) to perform 3D classification with 2 references (one good and one bad). Relion 3D auto-refine and post-processing then provided a 5 Å map with 438,001 selected particles. Those particles were imported into cryoSPARC again to perform non-uniform (NU) refinement^39^ and yielded the final reconstruction. Resolutions were estimated using the gold-standard Fourier shell correlation (GSFSC) with a 0.143 cut-off^40^. Local resolution was estimated using cryoSPARC^38^.

### Model building and refinement

The initial model of horse PepT1 was generated by AlphaFold2^41^. The TMD and ECD were individually docked into the cryo-EM density map in Chimera^42^. The map was sharpened in a convolutional neural network-based algorithm, DeepEMhancer^43^, with the highRes model. Processed map has significantly better resolvability of side chains and was used for model refinement. The docked model was manually adjusted in COOT^44^, and subjected to real-space refinement with secondary structure and geometry restraints in Phenix^45^. The EMRinger Score^46^ was calculated. All structure figures were prepared in ChimeraX^47^.

### Expression of horse PepT1 in HEK cells

The cDNA of horse PepT1 was cloned into a pEG BacMam vector with a C-terminal GFP tag. Mutations to horse PepT1 were generated using the QuikChange method (Stratagene) and the entire cDNA was sequenced to verify the mutation. The primers information is provided in Extended Data Table 2.

The HEK 293S cells in *FreeStyle 293* media (Invitrogen/Thermo Fisher) supplemented with 2% fetal bovine serum (FBS; Sigma) were maintained at 37°C with 8% CO_2_ in suspension culture at 100 rpm. Cells were plated one day before transfection to reach >90% confluency. Transfection of horse PepT1 plasmid or empty plasmid was performed with 293fectin transfection reagent (Invitrogen/Thermo Fisher) as per manufacturer’s instruction. Transfected cells were incubated at 37°C with 8% CO_2_ for 2 days. Cells on the plate were washed in PBS before scraping. Cell membranes were solubilized in the lysis buffer plus 1% LMNG and Protease Inhibitor Cocktail (Roche) for 1 h at 4°C. Insoluble fractions were pelleted by centrifugation and supernatants were run in SDS-PAGE. Bands of target proteins were visualized by western blotting with mouse anti-GFP (Invitrogen/Thermo Fisher) antibodies as primary antibodies and IRDye-800CW anti-mouse IgG (Licor) as secondary antibody. Images were taken on an Odyssey infrared scanner (Licor).

### Peptide uptake assay

The HEK 293S cells were plated on 6-well plates and transfected with the pEG BacMam plasmids with horse PepT1 as described above. Transfected cells were harvested after 2 days and transferred to 1.5 mL Eppendorf tubes. Cells were resuspended in transport buffer (20 mM HEPES, pH6.9, 140 mM NaCl, 2.5 mM KCl, 1.8 mM CaCl_2_, 1.0 mM MgCl_2_) shortly before assays. The ligand Ala-Ala were filtered through 0.45 µm nitrocellulose filters (Millipore). Cells were lysed in 0.1M NaOH and 1% SDS for 10 min. The radioactivity retained on the filters was determined by liquid scintillation counting. Counts per minute were converted to pmol by comparing to a standard curve plotted with known amounts of ^3^H-Ala-Ala. One-way ANOVA followed by the Dunnett’s post hoc test was performed. All statistical analyses were performed in GraphPad Prism 8.2.1.

### Proton transport assay

HEK293S cells from suspension culture were adjusted to a density of 0.75×10^6^ mL^-1^ with culture media. Transfection mixtures of the pEG BacMam plasmids with horse PepT1 were slowly added to cells under agitation. Transfected cells were aliquoted to poly-lysine-coated 96-well plates and incubated for 2 days. Cells were loaded with pHrodo Red dye (Invitrogen/Thermo Fisher) as per manufacturer’s instruction. Fluorescence was recorded in FlexStation 3 Multi-Mode Microplate Reader (Molecular Devices) at 37 °C. Shortly before the start of recording, cells on assay plates were exchanged to the pre-warmed transport buffer. The transport of peptide was initiated by adding 10 µL of ligands in the transport buffer to 90 µL of buffer in assay plates. Fluorescence readings at equilibrium (550 – 600 s) were averaged to represent intrasellar pH changes. Two-way ANOVA was used to compare the effect of outside pH. Two-tailed Student’s t-test was performed for comparison of mutants to the WT.

## Acknowledgments

This work was supported by grants from NIH (DK122784, HL086392, and GM098878 to M.Z.), Cancer Prevention and Research Institute of Texas (R1223 to M.Z.). We acknowledge the use of Princeton’s Imaging and Analysis Center, which is partially supported by the Princeton Center for Complex Materials, and the National Science Foundation (NSF)-MRSEC program (DMR-1420541).

## Data Availability

The atomic coordinate file of horse PepT1 in nanodisc has been deposited in the PDB (http://www.rcsb.org) under the accession codes 7S8U. The corresponding cryo-electron microscopy map has been deposited in the Electron Microscopy Data Bank (https://www.ebi.ac.uk/pdbe/emdb/) under the accession codes EMD-24922.

## Author Contributions

M.Z. conceived the project. J.S., M.H., Z.R., X.F., C.P., X.Y., and M.R. conducted experiments. J.S. and M.Z. wrote the paper.

---

**Extended Data Fig. 1:**
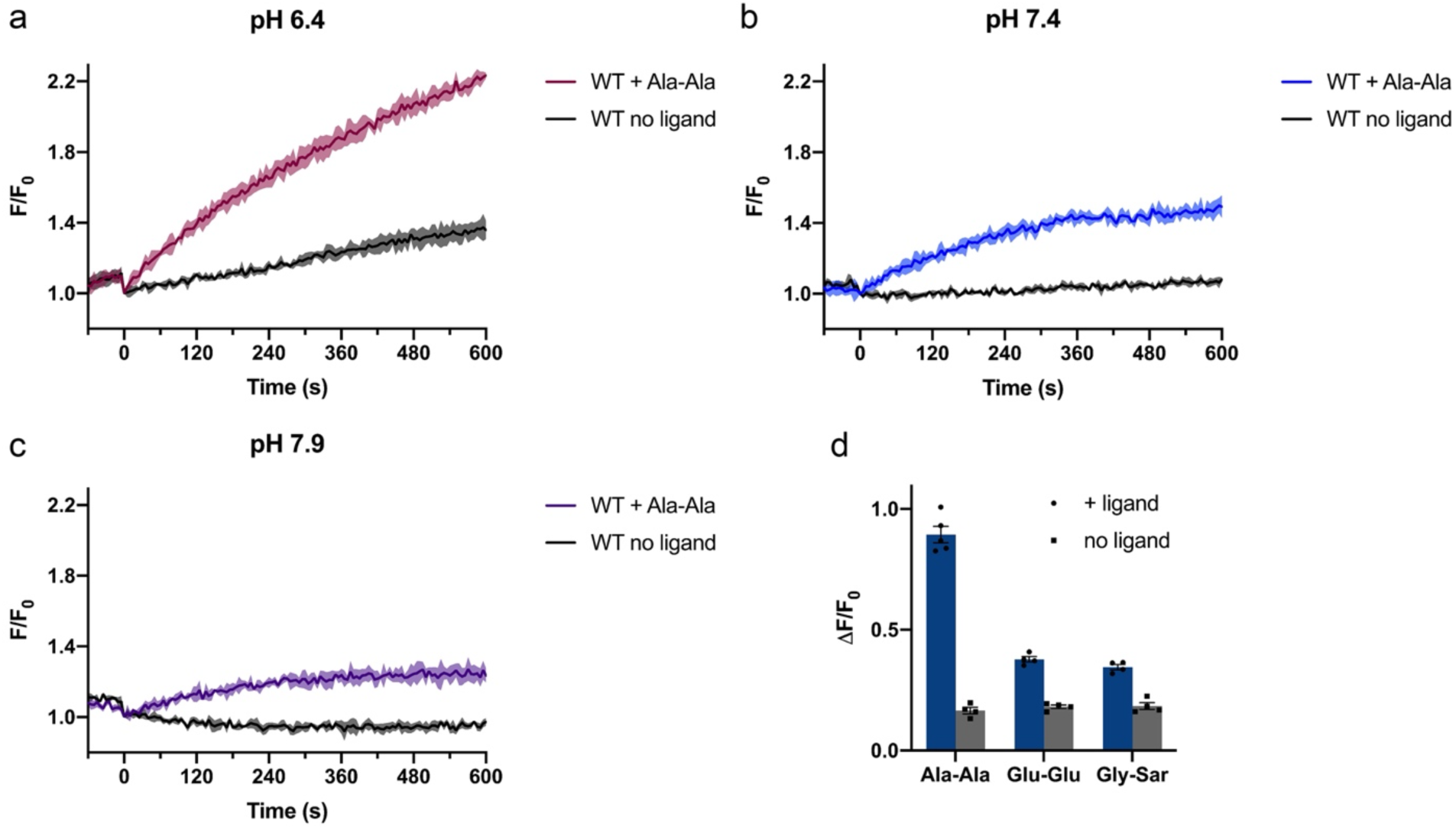
Time courses of pH changes in HEK293 cells induced by the peptide transport in the outside pH of 6.4 (**a**), 7.4 (**b**) and 7.9 (**c**). Solid lines represent the mean and shaded regions represent SD of four repeats. **d**, Cytosolic pH changes induced by different peptide ligands in the outside pH of 6.9.

**Extended Data Fig. 2:**
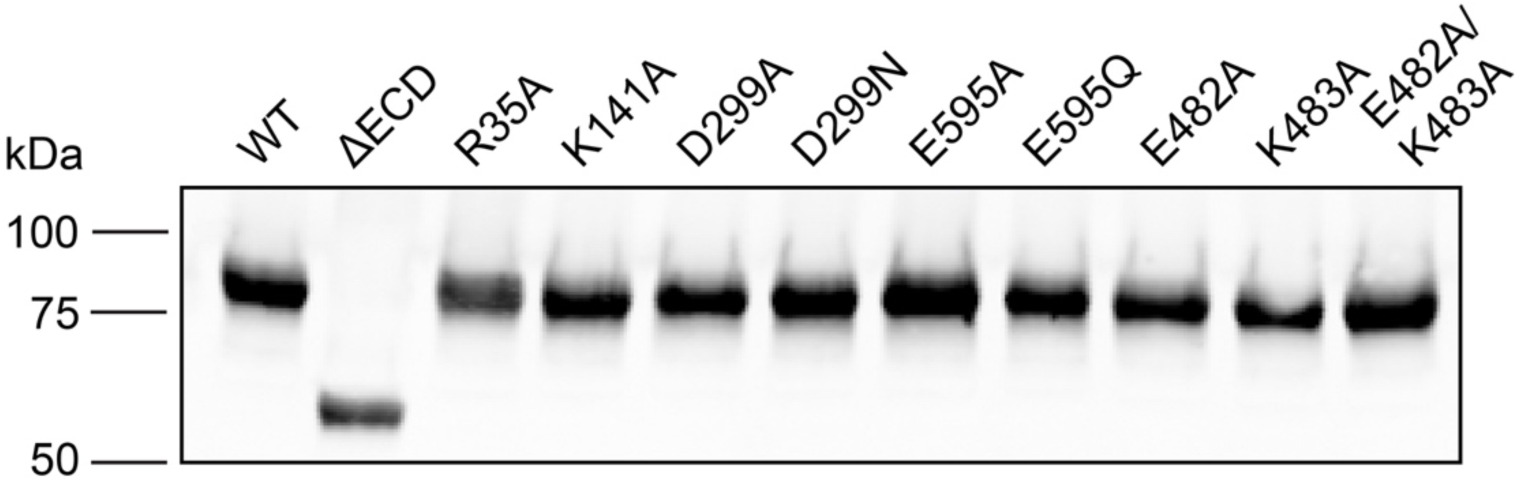
Assessment of expression level of GFP-tagged WT and mutant horse PepT1 in HEK293 cells by western blot.

**Extended Data Fig. 3:**
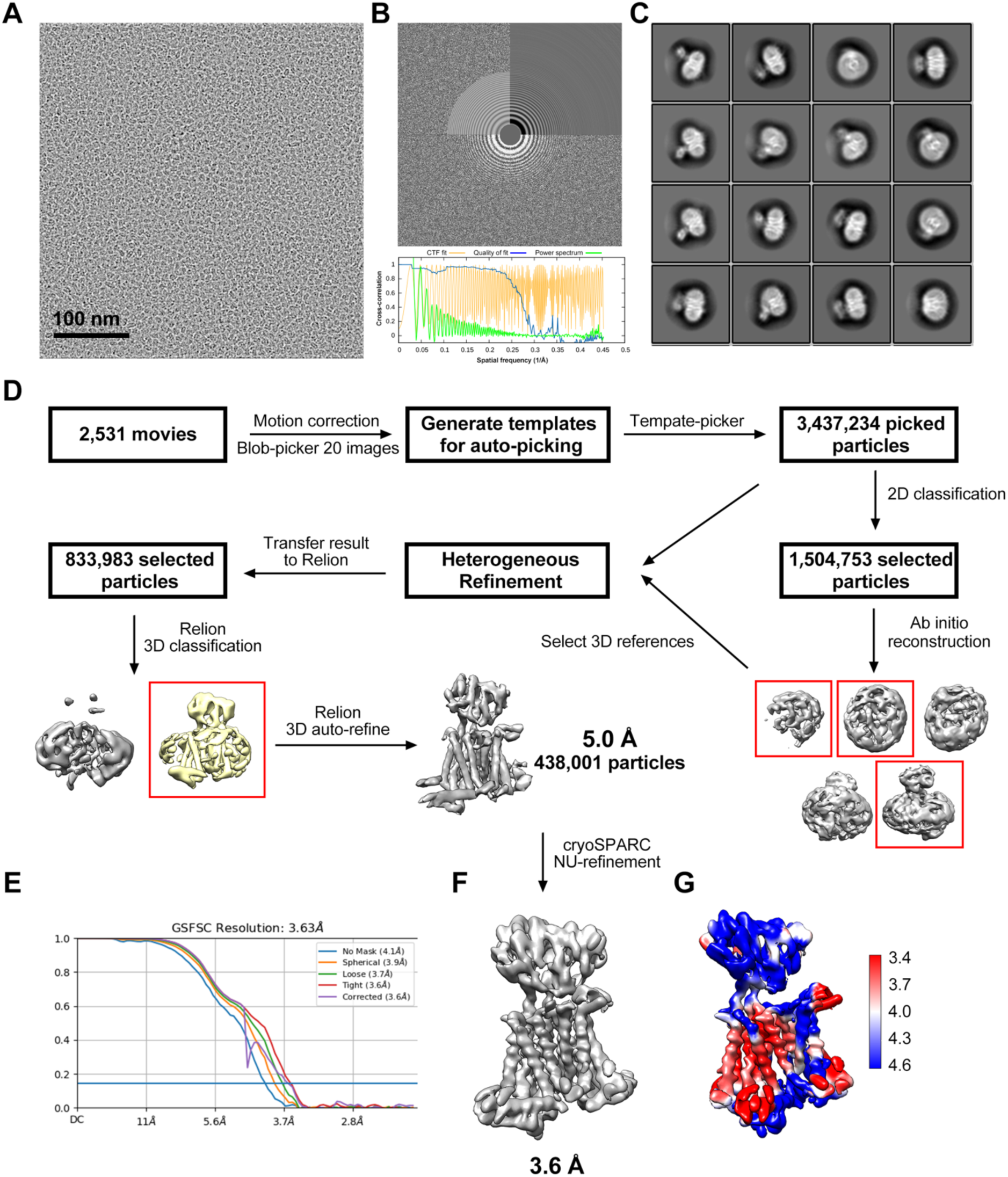
Cryo-EM data processing. **a**, A representative micrograph of horse PepT1, with **b**, its Fourier transform and contrast transfer function (CTF) fitting. **c**, Representative 2D class averages. **d**, The flow chart for data processing of PepT (methods). **e**, The gold-standard Fourier shell correlation (FSC) curve for the final map shown in **f**, and the local resolution map of horse PepT1 shown in **g**.

**Extended Data Fig. 4:**
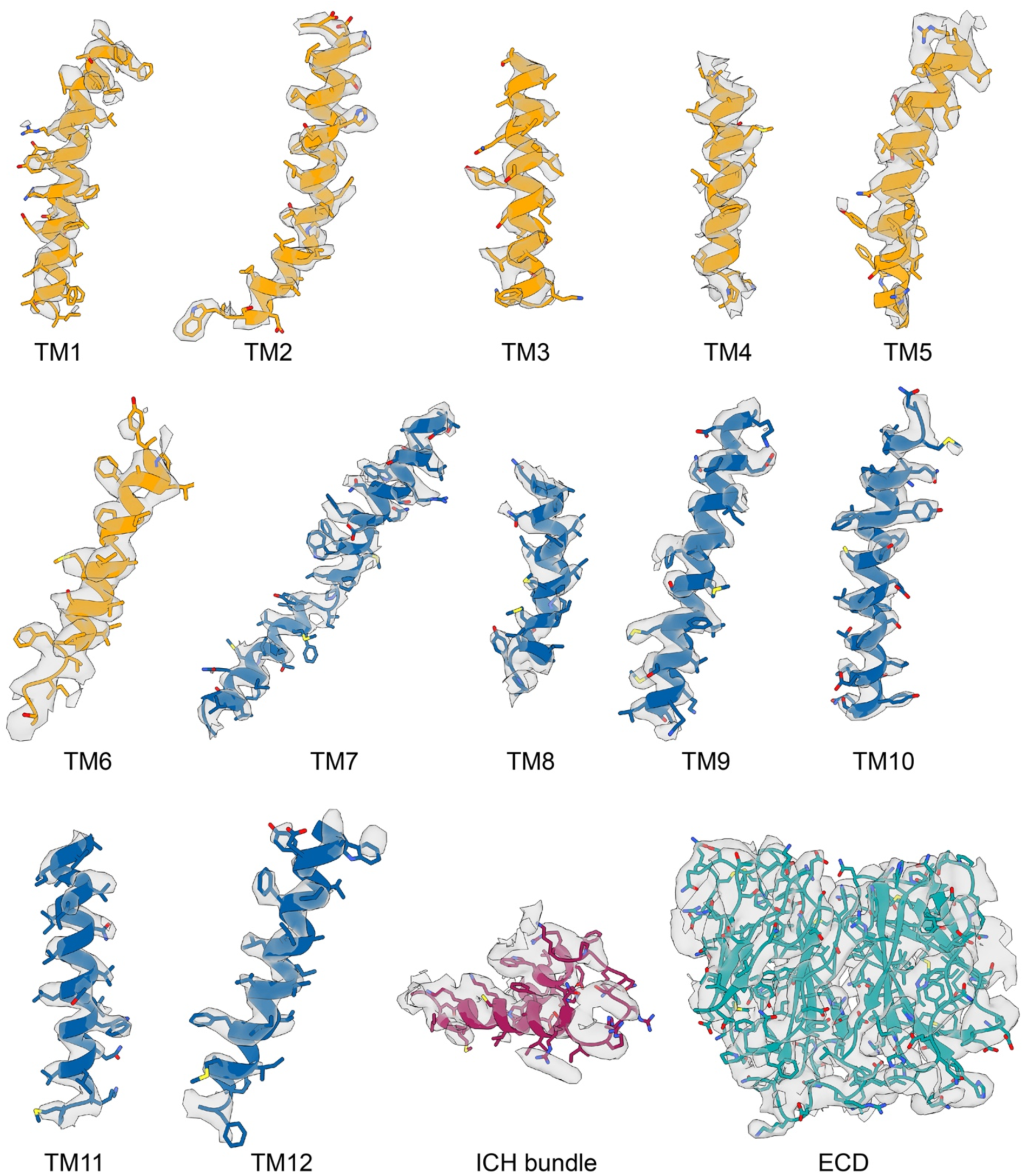
Cryo-EM map of TM helices, ICH bundle, and ECD of horse PepT1 in nanodisc.

**Extended Data Fig 5:**
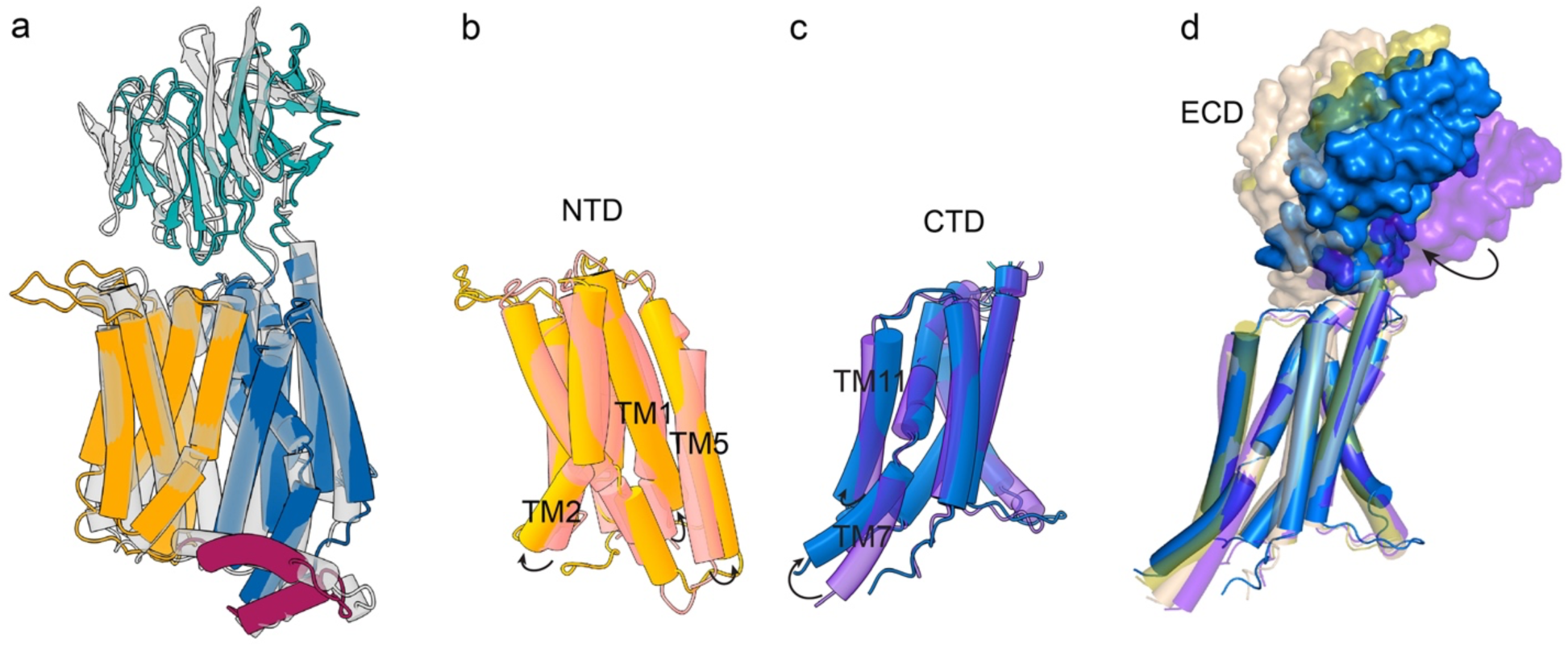
**a**, Structural comparison of horse PepT1 with the inward-facing human PepT2 (PDB ID: 7PMY) colored in translucent gray shows their overall structural similarity. Structural alignments of **b**, NTDs and **c**, CTDs of horse PepT1 and the outward-open rat PepT2 (PDB ID: 7NQK) colored in translucent pink and purple for NTD and CTD, respectively. The horse PepT1 are colored as in **Fig. 2c**. d, Variations in the relative position of ECD to the CTD among mammalian PepT structures. Blue, outward-facing horse PepT1 (this study); purple, outward-facing rat PepT2 (PDB ID: 7NQK); wheat, outward-facing human PepT1 (PDB ID: 7PMX); olive, inward-facing human PepT2 (PDB ID: 7PMY). Arrows indicate conformational differences of the aligned structures from horse PepT1.

**Extended Data Fig 6:**
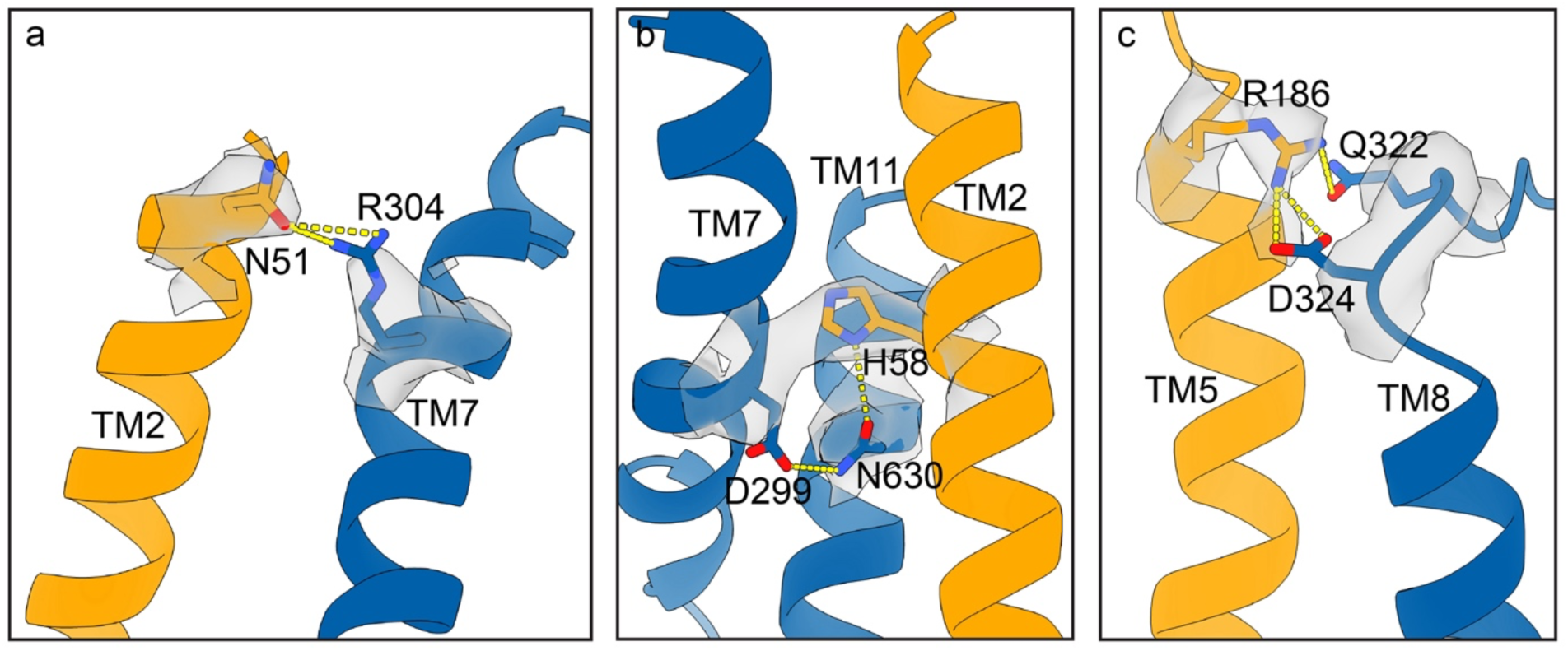
Hydrogen bonding and ionic interactions (yellow dash line) between NTD and CTD of horse PepT1. Electron density around the interacting residues is shown as translucent gray surface.

**Extended Data Fig 7:**
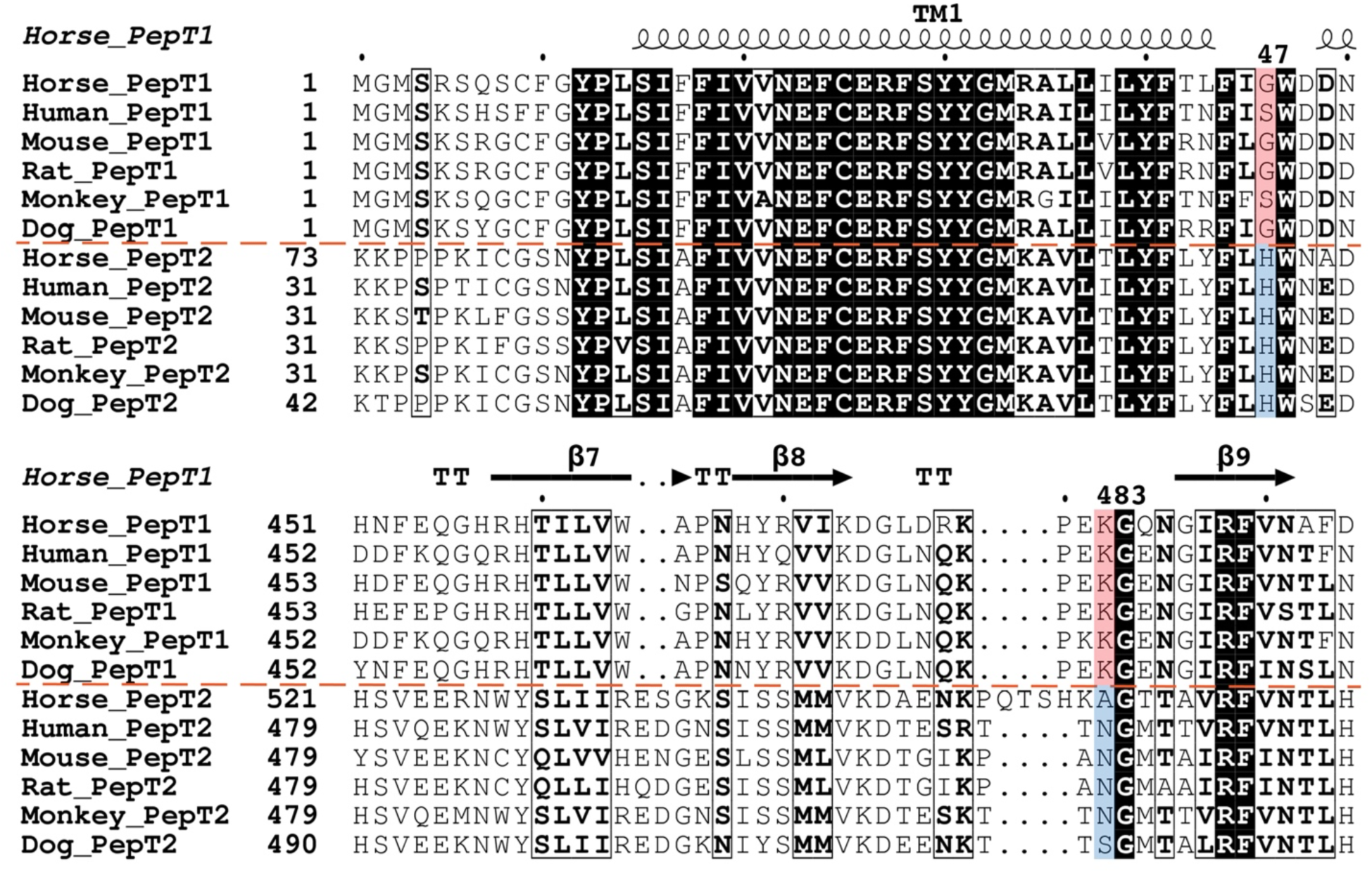
Sequence alignment of mammalian PepT1 and PepT2. PepT1 and PepT2 from horse (*Equus caballus*; UniProt ID: <u>F6SG69 and F6R282</u>), human (*Homo sapiens*; <u>P46059</u> and <u>Q16348</u>), mouse (*Mus musculus*; <u>Q9JIP7</u> and <u>Q9ES07</u>), rat (*Rattus norvegicus*; <u>P51574</u> and <u>Q63424</u>), monkey (*Macaca mulatta*; <u>F7H3Q3</u> and <u>Q6WFZ7</u>), and dog (*Canis familiaris*; <u>F1PTV0</u> and <u>E2QWX1</u>) are aligned using the Clustal Omega server. Secondary structural elements of horse PepT1 are marked above the alignment, and an orange dash lines separates the PepT1 and PepT2 group. Conserved residues in both PepT1 and PepT2 are indicated by black highlight and bold letter. Residue G47 and K483 of horse PepT1 and their equivalent residues are highlighted in red in the PepT1 group and in blue in the PepT2 group, respectively. The plot is prepared using the ESPript3 server.

**Extended Data Table 1.** Summary of Cryo-EM data collection, processing, and refinement.

| **Protein** | **Horse PepT1-MSP1D1 nanodisc** |
| --- | --- |
| **Cryo-EM Data Collection** |  |
| Voltage (kV) | 300 |
| Magnification (x) | 105,000 |
| Pixel Size (Å) | 1.114 |
| Electron exposure (e<sup>-</sup>/Å<sup>2</sup>/frame) | 1.56 |
| Defocus range (µm) | [-2.1, -1.9] |
| Number of image stacks | 2531 |
| Number of frames per stack | 32 |
| **Cryo-EM Data Processing** |  |
| Initial number of particles | 3,437,234 |
| Final number of particles | 438,001 |
| Symmetry imposed | C1 |
| Map sharpening B factor (Å<sup>2</sup>) | 150 |
| Map resolution (Å) | 3.6 |
| Map resolution range (Å) | 3.6 – 4.6 |
| FSC threshold | 0.143 |
| **Model Refinement** |  |
| Number of amino acids | 667 |
| Total non-hydrogen atoms | 5210 |
| Average B factor (Å<sup>2</sup>) | 122.76 |
| Bond length RMSD (Å) | 0.008 |
| Bond angle RMSD (°) | 1.177 |
| Ramachandran Plot |  |
| Favored (%) | 91.13 |
| Allowed (%) | 7.97 |
| Outliers (%) | 0.90 |
| Rotamer outliers (%) | 0.28 |
| EMRinger Score | 2.03 |

**Extended Data Table 2.**
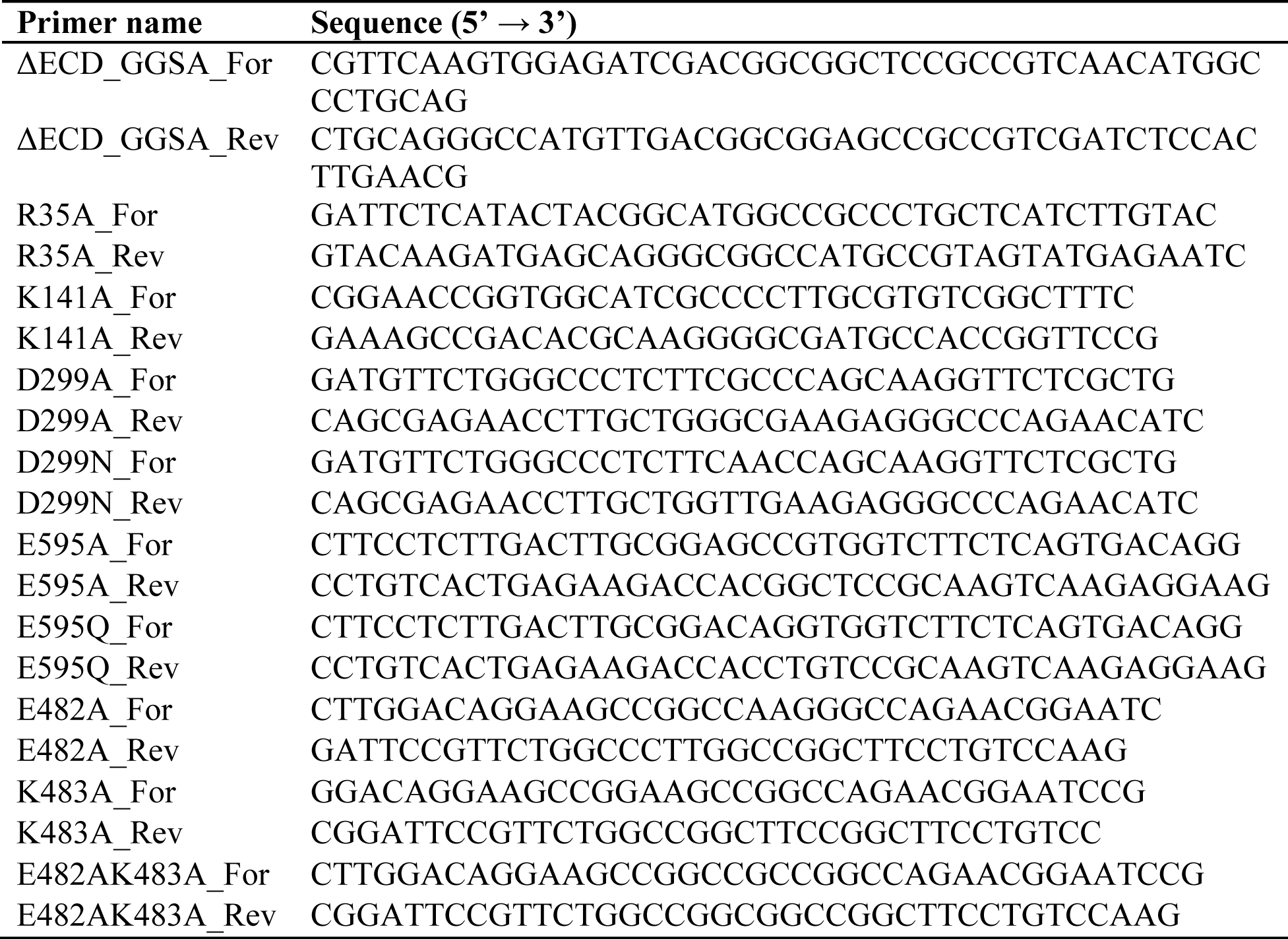
Primers used in this study.

## References

1 Daniel, H. & Kottra, G. The proton oligopeptide cotransporter family SLC15 in physiology and pharmacology. Pflugers Arch 447, 610–618, doi:10.1007/s00424-003-1101-4 (2004).

2 Qandeel, H. G., Duenes, J. A., Zheng, Y. & Sarr, M. G. Diurnal expression and function of peptide transporter 1 (PEPT1). J Surg Res 156, 123–128, doi:10.1016/j.jss.2009.03.052 (2009).

3 Matthews, D. M. Intestinal absorption of peptides. Biochem Soc Trans 11, 808–810, doi:10.1042/bst0110808 (1983).

4 Shen, H. et al. Localization of PEPT1 and PEPT2 proton-coupled oligopeptide transporter mRNA and protein in rat kidney. Am J Physiol 276, F658–665, doi:10.1152/ajprenal.1999.276.5.F658 (1999).

5 Kamal, M. A., Jiang, H., Hu, Y., Keep, R. F. & Smith, D. E. Influence of genetic knockout of Pept2 on the in vivo disposition of endogenous and exogenous carnosine in wild-type and Pept2 null mice. Am J Physiol Regul Integr Comp Physiol 296, R986–991, doi:10.1152/ajpregu.90744.2008 (2009).

6 Zhang, Y., Sun, J., Sun, Y., Wang, Y. & He, Z. Prodrug design targeting intestinal PepT1 for improved oral absorption: design and performance. Curr Drug Metab 14, 675–687, doi:10.2174/1389200211314060004 (2013).

7 Killer, M., Wald, J., Pieprzyk, J., Marlovits, T. C. & Low, C. Structural snapshots of human PepT1 and PepT2 reveal mechanistic insights into substrate and drug transport across epithelial membranes. Sci Adv 7, eabk3259, doi:10.1126/sciadv.abk3259 (2021).

8 Parker, J. L. et al. Cryo-EM structure of PepT2 reveals structural basis for proton-coupled peptide and prodrug transport in mammals. Sci Adv 7, doi:10.1126/sciadv.abh3355 (2021).

9 Newstead, S. et al. Crystal structure of a prokaryotic homologue of the mammalian oligopeptide-proton symporters, PepT1 and PepT2. EMBO J 30, 417–426, doi:10.1038/emboj.2010.309 (2011).

10 Guettou, F. et al. Structural insights into substrate recognition in proton-dependent oligopeptide transporters. EMBO Rep 14, 804–810, doi:10.1038/embor.2013.107 (2013).

11 Doki, S. et al. Structural basis for dynamic mechanism of proton-coupled symport by the peptide transporter POT. Proc Natl Acad Sci U S A 110, 11343–11348, doi:10.1073/pnas.1301079110 (2013).

12 Lyons, J. A. et al. Structural basis for polyspecificity in the POT family of proton-coupled oligopeptide transporters. EMBO Rep 15, 886–893, doi:10.15252/embr.201338403 (2014).

13 Guettou, F. et al. Selectivity mechanism of a bacterial homolog of the human drug-peptide transporters PepT1 and PepT2. Nat Struct Mol Biol 21, 728–731, doi:10.1038/nsmb.2860 (2014).

14 Boggavarapu, R., Jeckelmann, J. M., Harder, D., Ucurum, Z. & Fotiadis, D. Role of electrostatic interactions for ligand recognition and specificity of peptide transporters. BMC Biol 13, 58, doi:10.1186/s12915-015-0167-8 (2015).

15 Parker, J. L. et al. Proton movement and coupling in the POT family of peptide transporters. Proc Natl Acad Sci U S A 114, 13182–13187, doi:10.1073/pnas.1710727114 (2017).

16 Quistgaard, E. M., Martinez Molledo, M. & Low, C. Structure determination of a major facilitator peptide transporter: Inward facing PepTSt from Streptococcus thermophiles crystallized in space group P3121. PLoS One 12, e0173126, doi:10.1371/journal.pone.0173126 (2017).

17 Martinez Molledo, M., Quistgaard, E. M. & Low, C. Tripeptide binding in a proton-dependent oligopeptide transporter. FEBS Lett 592, 3239–3247, doi:10.1002/1873-3468.13246 (2018).

18 Martinez Molledo, M., Quistgaard, E. M., Flayhan, A., Pieprzyk, J. & Low, C. Multispecific Substrate Recognition in a Proton-Dependent Oligopeptide Transporter. Structure 26, 467–476 e464, doi:10.1016/j.str.2018.01.005 (2018).

19 Minhas, G. S. & Newstead, S. Structural basis for prodrug recognition by the SLC15 family of proton-coupled peptide transporters. Proc Natl Acad Sci U S A 116, 804–809, doi:10.1073/pnas.1813715116 (2019).

20 Ural-Blimke, Y. et al. Structure of Prototypic Peptide Transporter DtpA from E. coli in Complex with Valganciclovir Provides Insights into Drug Binding of Human PepT1. J Am Chem Soc 141, 2404–2412, doi:10.1021/jacs.8b11343 (2019).

21 Newstead, S. Recent advances in understanding proton coupled peptide transport via the POT family. Curr Opin Struct Biol 45, 17–24, doi:10.1016/j.sbi.2016.10.018 (2017).

22 Drew, D., North, R. A., Nagarathinam, K. & Tanabe, M. Structures and General Transport Mechanisms by the Major Facilitator Superfamily (MFS). Chem Rev 121, 5289–5335, doi:10.1021/acs.chemrev.0c00983 (2021).

23 Beale, J. H. et al. Crystal Structures of the Extracellular Domain from PepT1 and PepT2 Provide Novel Insights into Mammalian Peptide Transport. Structure 23, 1889–1899, doi:10.1016/j.str.2015.07.016 (2015).

24 Steel, A. et al. Stoichiometry and pH dependence of the rabbit proton-dependent oligopeptide transporter PepT1. J Physiol 498 (Pt 3), 563–569, doi:10.1113/jphysiol.1997.sp021883 (1997).

25 Terada, T., Irie, M., Okuda, M. & Inui, K. Genetic variant Arg57His in human H+/peptide cotransporter 2 causes a complete loss of transport function. Biochem Biophys Res Commun 316, 416–420, doi:10.1016/j.bbrc.2004.02.063 (2004).

26 Xu, L., Haworth, I. S., Kulkarni, A. A., Bolger, M. B. & Davies, D. L. Mutagenesis and cysteine scanning of transmembrane domain 10 of the human dipeptide transporter. Pharm Res 26, 2358–2366, doi:10.1007/s11095-009-9952-9 (2009).

27 Ramamoorthy, S. et al. Proton/peptide cotransporter (PEPT 2) from human kidney: functional characterization and chromosomal localization. Biochim Biophys Acta 1240, 1–4, doi:10.1016/0005-2736(95)00178-7 (1995).

28 Chen, X. Z., Zhu, T., Smith, D. E. & Hediger, M. A. Stoichiometry and kinetics of the high-affinity H+-coupled peptide transporter PepT2. J Biol Chem 274, 2773–2779, doi:10.1074/jbc.274.5.2773 (1999).

29 Martens, C. et al. Lipids modulate the conformational dynamics of a secondary multidrug transporter. Nature structural & molecular biology 23, 744–751, doi:10.1038/nsmb.3262 (2016).

30 Autzen, H. E. et al. Structure of the human TRPM4 ion channel in a lipid nanodisc. Science (New York, N.Y.) 359, 228–232, doi:10.1126/science.aar4510 (2018).

31 Pan, Y. et al. Structural basis of ion transport and inhibition in ferroportin. Nat Commun 11, 5686, doi:10.1038/s41467-020-19458-6 (2020).

32 Mastronarde, D. N. Automated electron microscope tomography using robust prediction of specimen movements. J Struct Biol 152, 36–51, doi:10.1016/j.jsb.2005.07.007 (2005).

33 Zheng, S. Q. et al. MotionCor2: anisotropic correction of beam-induced motion for improved cryo-electron microscopy. Nat Methods 14, 331–332, doi:10.1038/nmeth.4193 (2017).

34 Grant, T. & Grigorieff, N. Measuring the optimal exposure for single particle cryo-EM using a 2.6 A reconstruction of rotavirus VP6. Elife 4, e06980, doi:10.7554/eLife.06980 (2015).

35 Scheres, S. H. Semi-automated selection of cryo-EM particles in RELION-1.3. J Struct Biol 189, 114–122, doi:10.1016/j.jsb.2014.11.010 (2015).

36 Scheres, S. H. RELION: implementation of a Bayesian approach to cryo-EM structure determination. J Struct Biol 180, 519–530, doi:10.1016/j.jsb.2012.09.006 (2012).

37 Kimanius, D., Forsberg, B. O., Scheres, S. H. & Lindahl, E. Accelerated cryo-EM structure determination with parallelisation using GPUs in RELION-2. Elife 5, doi:10.7554/eLife.18722 (2016).

38 Punjani, A., Rubinstein, J. L., Fleet, D. J. & Brubaker, M. A. cryoSPARC: algorithms for rapid unsupervised cryo-EM structure determination. Nat Methods 14, 290–296, doi:10.1038/nmeth.4169 (2017).

39 Punjani, A., Zhang, H. & Fleet, D. J. Non-uniform refinement: adaptive regularization improves single-particle cryo-EM reconstruction. Nat Methods 17, 1214–1221, doi:10.1038/s41592-020-00990-8 (2020).

40 Rosenthal, P. B. & Henderson, R. Optimal determination of particle orientation, absolute hand, and contrast loss in single-particle electron cryomicroscopy. J Mol Biol 333, 721–745 (2003).

41 Jumper, J. et al. Highly accurate protein structure prediction with AlphaFold. Nature 596, 583–589, doi:10.1038/s41586-021-03819-2 (2021).

42 Pettersen, E. F. et al. UCSF Chimera--a visualization system for exploratory research and analysis. Journal of computational chemistry 25, 1605–1612, doi:10.1002/jcc.20084 (2004).

43 Sanchez-Garcia, R. et al. DeepEMhancer: a deep learning solution for cryo-EM volume post-processing. Commun Biol 4, 874, doi:10.1038/s42003-021-02399-1 (2021).

44 Emsley, P., Lohkamp, B., Scott, W. G. & Cowtan, K. Features and development of Coot. Acta crystallographica. Section D, Biological crystallography 66, 486–501, doi:10.1107/S0907444910007493 (2010).

45 Adams, P. D. et al. PHENIX: a comprehensive Python-based system for macromolecular structure solution. Acta crystallographica. Section D, Biological crystallography 66, 213–221, doi:10.1107/S0907444909052925 (2010).

46 Barad, B. A. et al. EMRinger: side chain-directed model and map validation for 3D cryo-electron microscopy. Nature methods 12, 943–946, doi:10.1038/nmeth.3541 (2015).

47 Pettersen, E. F. et al. UCSF ChimeraX: Structure visualization for researchers, educators, and developers. Protein Sci 30, 70–82, doi:10.1002/pro.3943 (2021).

